# Modulation of Arm Swing Frequency and Gait Using Rhythmic Tactile Feedback

**DOI:** 10.1101/2022.06.03.494736

**Authors:** Mohsen Alizadeh Noghani, Md. Tanzid Hossain, Babak Hejrati

**Affiliations:** Biorobotics and Biomechanics Lab, University of Maine, Orono, ME 04469 USA

**Keywords:** Haptic feedback, rhythmic stimuli, interlimb coupling, arm swing, gait rehabilitation, dynamic system response

## Abstract

Due to the neural coupling between upper and lower limbs and the importance of interlimb coordination in human gait, it has been recommended that focusing on appropriate arm swing should be a part of gait rehabilitation in individuals with walking impairments. Despite its vital importance, there is a lack of effective methods to exploit the potential of arm swing inclusion for gait rehabilitation. In this work, we present a lightweight and wireless haptic feedback system that provides highly synchronized vibrotactile cues to the arms to manipulate arm swing and investigate the effects of this manipulation on the subjects’ gait in a study with 12 participants (20-44 years). We found the developed system effectively adjusted the subjects’ arm swing and stride cycle times by significantly reducing and increasing those parameters by up to 20% and 35%, respectively, compared to their baseline values during normal walking with no feedback. Particularly, the reduction of arms’ and legs’ cycle times translated into a substantial increase of about 19.3% (on average) in walking speed. The response of the subjects to the feedback was also quantified in both transient and steady-state walking. The analysis of settling times from the transient responses revealed a fast and similar adaptation of both arms’ and legs’ movements to the feedback for reducing cycle time (i.e., increasing speed). Conversely, larger settling times and the time differences between arms’ and legs’ responses were observed due to feedback for increasing cycle times (i.e., reducing speed). The results clearly demonstrate the potential of the developed system to induce different arm-swing patterns as well as the ability of the proposed method to modulate key gait parameters through capitalizing on the interlimb neural coupling with implications for gait training.

## I. Introduction

The role of arm swing in human gait has been the subject of extensive research and discussion, and although no unified theory has been proposed for its function, reducing energy expenditure is well-established as one of its effects [1], [2]. Importantly, evidence suggests the existence of neural couplings between the upper and lower limbs and points to the crucial role of appropriate arm swing in interlimb coordination and improving central pattern generators’ (CPG) function and training. For instance, by analyzing EEG signals of the Supplementary Motor Area (SMA) of the cortex, which is associated with coordinating the motor system, Weersink et al. [3] found that preventing arm swing decreases brain activity in that region. Bloom and Hejrati [4] demonstrated that constraining the forearms significantly impacts the patterns of arm swing and interlimb coordination during walking, while increasing the muscle activities of the biceps, trapeziuses, and posterior deltoids that controlled the shoulder motion and trunk rotation. Another study focusing on the timing of EMG signals of the upper and lower limb muscles revealed that the arm activity drives (leads) the lower limbs’ [5]. A similar effect was observed by studying gait initiation in older adults which found that both backward and forward swings of the arms drive the upper leg muscles’ activities [6]. La Scaleia et al. [7] predicted lower limb kinematics (e.g., the thigh, shank, and foot angles) from the EMG signals of the deltoid (shoulder) muscles using a template-matching method. In a study where the participants’ legs were suspended, in the majority of subjects, rhythmic motion of the arms in the antero-posterior direction elicited muscle activity and movement patterns in the legs similar to those created during walking [8]. Hejrati et al. [9] created a data-driven linear time-invariant transfer function to predict the correct arm swing based on the realtime measurements of thighs’ angular velocity. The growing evidence for the role of arms in human locomotion has resulted in recommendations for their inclusion in rehabilitation or gait training of those with disrupted coordination abilities, such as stroke or Parkinson’s disease (PD) patients [10], [11].

Several studies in recent years have investigated the effect of adjusted arm swing on movements of clinical populations and older adults. In a study of traumatic brain injury (TBI) patients by Ustinova et al. [12], they received instructions to increase their arm swing amplitude and synchronization, which improved shoulder and hip motion coupling of the subjects and increased their step lengths. Other works have found that a higher arm swing amplitude leads to improved trunk stability in young adults [13] and the elderly [14]. In a study of PD patients by Weersink et al. [15], it was observed that when the subjects walked with an increased arm swing range of motion, their step length, walking speed, and brain activity patterns resembled those of healthy controls. In another study of PD patients and also older adults, instructions to increase arm swing amplitude and frequency resulted in improved stride velocities in both groups [16]. A 5-weeks training course of only arm cycling in a stationary position for chronic stroke patients designed by Kaupp et al. [17] resulted in improvements in interlimb coupling and clinically significant enhancements of balance and gait. It should be noted that increases in stride velocity are linked to a reduced rate of mortality in older adults [18]. Therefore, healthy older adults could benefit from interventions that could improve their walking speed as well.

The studies mentioned above provided verbal instructions or feedback to the users; while that may be effective for in-person clinical sessions, biofeedback systems, which provide auditory, visual, or tactile stimuli, can enable intervention or training in home-based settings [19]. Among these 3 modes, auditory feedback in the form of rhythmic auditory stimulation (RAS) [20], where the user should move in sync with sound cues (generated by a metronome in the simplest case) or music, is the most common form. Training with RAS can improve spatiotemporal gait parameters (e.g., cadence, stride length, and stride velocity) and functional mobility measures such as timed up and go as well as arm function in stroke [21], [22] and PD [23] patients. In a treadmill walking study, Ford et al. [24] instructed hemiparetic post-stroke patients to swing their arms and legs to a metronome frequency of about 1.8 Hz which led to higher arm swing range of motion (ROM) and stride length [24]. Mainka et al. [25] designed a system that adjusted the rhythm of the music based on the acceleration of the subject’s lower arm. They demonstrated that when PD patients were instructed to swing their arms according to the musical rhythm, parameters such as arm swing ROM and regularity, cadence, and stride velocity increased compared to their normal walking.

Despite most studies utilizing the auditory mode, research supports the idea that tactile feedback may be better suited for providing adjustment signals than both visual and auditory cues. It has been noted that tactile feedback can be presented more privately and could be easier to perceive in noisy or visually cluttered environments [26]; in such scenarios, awareness of external visual and auditory cues in the environment is essential for safe navigation. For example, sounds from the environment improve balance while walking [27] which would be blocked when wearing headphones to listen to auditory cues. Therefore, in accordance with multiple-resource theory [28], it is recommended to relay information across other sensory channels such as tactile. Pitts and Sarter [29] observed that in both young and older adults, response to tactile stimuli is faster than those from visual and auditory sources; similar results, along with a higher detection rate of tactile modality were reported by Bajpai et al. [30]. Sklar and Sarter [31] found that tactile cues had higher detection rates and faster response times, and did not interfere with nor were affected by visual feedback. Comparing auditory and tactile warnings in noisy environments, Murata et al. [32] reported tactile cues to perform as well or better than auditory sources. Tactile cues were found to be superior in terms of reaction time to auditory signals in a multitasking scenario such as a complex phone conversation while driving [33]. Additionally, a tactile feedback system is a practical option to be conveniently used in real-world indoor and outdoor settings such as home and community rather than the other systems that are mainly bound to research and clinical settings.

Therefore, utilizing tactile feedback for motor training, including for arm swing, has attracted increasing attention. In a study of PD patients, Thompson et al. [34] used vibrotactile cues on the arm to provide feedback for increasing arm swing amplitude which resulted in increases in that parameter, as well as longer steps and lower walking cadence but no significant change in walking speed. Lee et al. [35] also targeted arm swing range of motion, but they provided cues for both forward and backward swing ranges of motion. The healthy subjects achieved the required increases in arm swing angle which led to higher values of stride length while the changes in gait speed did not reach a significant level. In additional testing with one hemiparetic stroke patient, improvements in symmetry ratio were observed. In both subject groups, providing feedback to one arm influenced the range of motion of the other arm as well. In a study of PD patients by Kishi et al. [36], a wearable robot with a mass of 4.6 kg applied rhythmic kinesthetic haptic feedback to the arms with the goal of increasing their backward swing. Providing the feedback increased arm swing amplitude and improved stride length, stride velocity, and gait variability. Comparing the subject’s gait in pre-intervention and post-intervention trials showed a lasting effect for those improvements immediately after the training session.

While the research studies above have shown promise for increasing arm swing amplitude, the effects of providing rhythmic tactile feedback to adjust the arm swing cycle time (i.e., the reciprocal of the frequency) is unexplored. Notably, previous research has shown that the arm swing frequency is directly and highly correlated with its amplitude and also walking speed and cadence [37]. In this work, we (1) present a lightweight, wearable, and wireless feedback system capable of providing synchronized vibrotactile feedback to the arms. Further, by providing the feedback at various rhythms and in a variety of conditions in a study with 12 participants, we quantify its effect on the subjects’ (2) arm swing amplitude and cycle time, (3) foot cycle time, stride length, and stride velocity. Finally, we present (4) the participants’ subjective ratings and preferences which evaluated various characteristics of the system.

## II. Methods

### A. Feedback System

The system was a modified version of the wireless and modular system developed by Alizadeh Noghani et al. [38], [39], which uses a smartphone as the main controller. As shown in Fig. 1a, vibrotactors (10 mm in diameter) were placed on the anterior and posterior sides of the arms, and the electronics units, each consisting of an ESP8266 micro-controller, a battery, and a driving circuit for the vibrotactors, as well as the arm IMUs (Xsens Technologies B.V., Enschede, The Netherlands) were attached on their lateral sides. The vibrotactors were attached using a Micropore tape and kinesiology tape (not shown in the figure) was used on the top of to ensure that the vibrotactors maintained their contact with the skin. To measure spatiotemporal parameters, foot IMUs were attached to the posterior side of the subjects’ shoes. The system’s components added a mass of 58g to each arm making the system lightweight and comfortable to use.

**Fig. 1:**
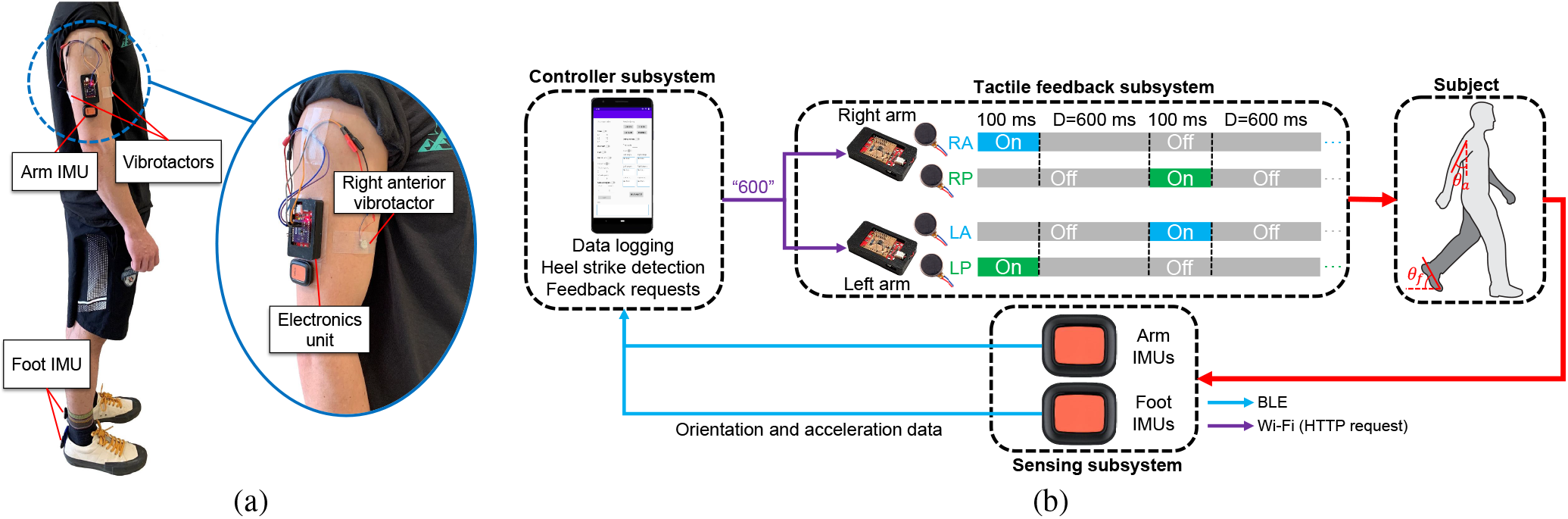
(a) The system worn by a subject and (b) the components of the feedback system with the feedback method shown for one cycle, in which the vibrotactors on the anterior side and posterior side of opposite arms turned on together (RA with LP and RP with LA) and there was a delay of *D* between the activations of the same arm’s anterior and posterior vibrotactors.

The IMUs streamed orientation and acceleration data at 60 Hz to the phone via Bluetooth Low Energy (BLE); the data of all sensors were logged in CSV files, and, in order to count the number of gait cycles, the data of the right foot IMU was processed in real-time to obtain the sagittal angle of the foot (*θ*_*f*_) and thus heel strike events (Fig. 2). The ESP8266 microcontrollers were programmed as HTTP servers connected to a WiFi network created by the phone and received requests from the Android application containing information about the time delay *D* between turning off one vibrotactor and turning on the other.

**Fig. 2:**
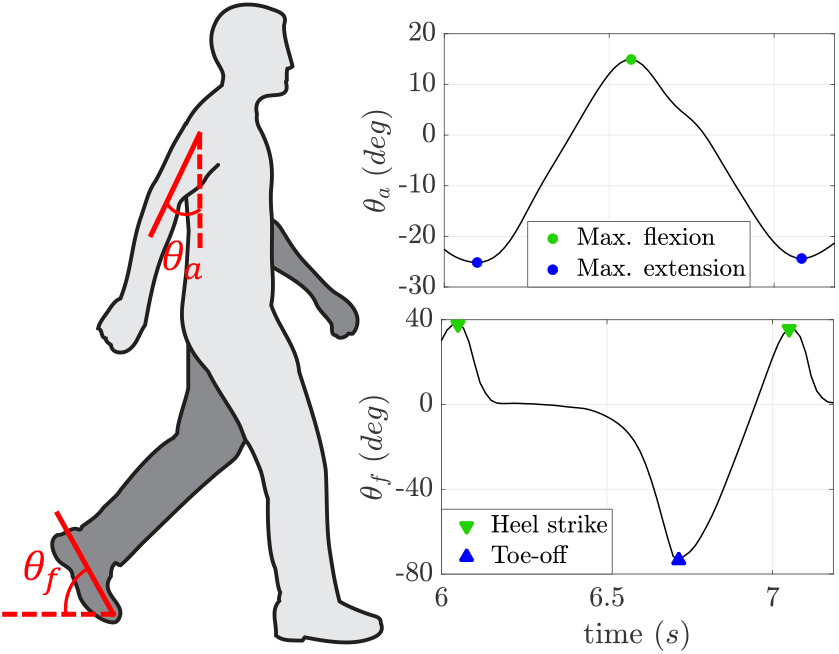
An example of the sagittal arm and foot angles of the same side as obtained from the IMU data, shown for one gait cycle.

As shown in Fig. 1b for one feedback cycle, to convey information about the rhythm of arm swing a synchronized and “diagonal” feedback pattern was employed, where the right anterior (RA) and the left posterior (LP) vibrotactors were activated at the same time and, the right posterior (RP) and left anterior (LA) vibrotactors were activated in a similar manner. During the experiments, the subjects were asked to adjust their arm swing so that the peak of the forward swing (i.e., maximum flexion in Fig. 2) occurred at the same time as the vibration on the anterior side, and the peak of the backward swing (i.e., maximum extension in Fig. 2) at the same time as the vibration on the posterior side. For this diagonal feedback scheme to be effective and usable, the activation of the vibrotactors should be (1) comfortable, (2) noticeable, (3) short enough to be perceived as happening for an instant as they are supposed to indicate only the maxima or minima of the arm angle (i.e., peak angle events), and (4) synchronized (i.e., each pair of the RA-LP and RP-LA turned on at the same time). Next, we discuss how we tried to satisfy the latter 3 requirements. Additionally, as will be discussed in Section II-A, we collected subjective ratings on the 4 criteria after the experiments to evaluate how the subjects perceived the system.

The used vibrotactors were brushed DC motors and, therefore, they exhibited the response of a first-order system. When the pin connected to a vibrotactor was set high, the motor reached its operational speed and, thus, its highest vibration intensity only after its settling time of about 158 ms. Therefore, turning off the motor too quickly would result in a low vibration intensity and difficulty in perceiving the tactile cue. On the other hand, the vibration had to be short enough to be associated with the peak angle events by the subject. In pilot trials, different durations for the on-time of the pins (*t*_*on*_) were tested and a value of 100 ms for *t*_*on*_ was found to attain a good balance between the duration of vibration and perceptibility, thus, addressing criteria (2) and (3). Analyzing the acceleration envelope profile of the vibrotactor during its on-time showed that it reached 56% percent of the maximum attainable acceleration at its peak. Further, the acceleration was at or higher than 90% of the peak value for 24 ms.

To achieve synchronization between the ESP8266 microcontrollers, the Android application and the HTTP server code on the ESP8266 microcontrollers were tuned for maximum determinism and minimum delay between the activation of the vibrotactors, where the delay, *D*, was defined as the time difference between the rising edges of the pairs of pins that the vibrotactors of one arm (i.e., the anterior and posterior) were connected to. Using 1000 measurements in operational conditions, the delay was found to be 1.5 ± 1.7 ms (mean ± standard deviation) with a worst-case delay of 17.7 ms and 95 and 99 percentile values of 3.9 ms and 9.6 ms, respectively. These results demonstrate that the system achieved a small delay and a high degree of determinism and, therefore, satisfied criterion (4).

### B. Experiment Design

After placing the system on the subjects, they were asked to perform the following trials on a 54 m *×* 3 m walkway,

- Fast walking (*F*): 1 lap. The subjects were asked to walk at their comfortably fast speed.
- Normal walking (*N*): 2 laps. Starting from the 11th stride, the next consecutive 100 right strides were used to calculate the average cycle time of the right foot 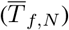.

The feedback trials were performed next. The subjects were asked to start walking at their normal speed and adjust their arm swing to match the vibrations on their arms according to the diagonal feedback scheme discussed in Section II-A. This “open-loop” method, where the information from the subject’s performance during their walking is not used to adjust the vibration pattern or timing, was selected because of its success in a previous work that used vibrotactile feedback to regulate the timing of heel strikes [40]. To minimize the possibility of receiving auditory cues from the vibrotactors, the subjects put on a headphone that played white noise during the trials. For each arm, the delay *D* was calculated by multiplying a coefficient, *k*, by half of the gait cycle time calculated from trial *N* 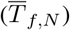 as shown in Eq. 1. We used the foot cycle time as equivalent to arm cycle time due to the 1:1 arm-to-leg swing frequency ratio [37] and the fact that finding the peak of foot angle in real-time was simpler as it is more regular and predictable during a gait cycle.

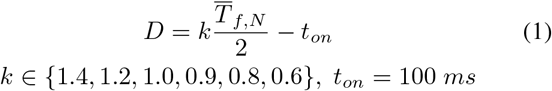

The time between successive activations of a vibrotactor was 2*D* + 2*t*_*on*_ (see Fig. 1b). We call that the feedback cycle time (*T*_*F B*_) and by using Eq. 1 we have,

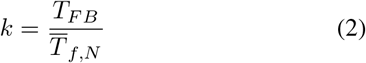

In other words, *k* specified the desired arm cycle time of the feedback trial relative to the stride time of normal walking. Here *k* ≥ 1 indicates desired arm cycles longer than or equal to the normal stride time and *k* < 1 results in shorter arm cycles. For each trial, the value of *D* was calculated in the Android application and, after the detection of the 10th heel strike, was sent to the electronics units in HTTP requests automatically. For example, using Eq. 1, for an average stride time of 1000 ms in normal walking, *k* = 1.4 would result in to *D* = 600 ms.

After the normal walking trial, two feedback familiarization trials (1 lap each, *k* = 1.45 and *k* = 0.95) were performed. With the exception of the two familiarization trials and trial *HI*_0.8_, no information about whether the feedback rhythm would be faster or slower than normal or when it would start was provided. The feedback trials are introduced in the list below:

- *H*_*x*_ (6 trials, 1 lap each): where *x* is the value of *k* in the trial.
- *HM*_0.9_ (2 laps): feedback was enabled for 40 cycles with *k* = 0.9. Then, it was disabled for 40 cycles and the subject was asked to maintain the same arm swing rhythm. The goal was to investigate whether the subjects could maintain the same gait after a short period of receiving feedback.
- *HO*_0.9_ (1 lap): feedback was provided to only one arm (*k* = 0.9), with half the subjects receiving it on the dominant and the other half on the non-dominant side. The subjects were instructed to maintain symmetry between the two sides. With this trial, we aimed to study the differences (if any) between the side that received feedback and the side that did not, as well as any changes in gait compared to *H*_0.9_.
- *HI*_0.8_ (1 lap): in this trial with *k* = 0.8, the subjects were told that the rhythm of the feedback would be 40% faster than their normal walking. This is the only trial in which the subjects received any information about the feedback. The goal was to see whether providing this information, which exaggerated how fast the timing of the feedback would be, resulted in any differences in the subjects’ gaits compared to *H*_0.8_, which corresponded to a feedback rhythm of 20% faster than normal walking cycle time.

Regarding the order of the feedback trials, the overall sequence was randomized and different for each subject, with the following exceptions: (1) *H*_0.9_ and *HO*_0.9_ were performed consecutively, but their order was different for each subject, (2) to reduce the possibility of biasing the subject’s gait, the three trials with the same value of *k* (*H*_0.9_, *HM*_0.9_, and *HO*_0.9_) were never performed consecutively.

After completing trials *H*_0.9_ and *HO*_0.9_, the subjects were asked whether feedback on one side or both sides was easier to follow, or if there was no difference between them. Additionally, immediately after the end of the experiments, they filled out a questionnaire rating their experience with the system on a scale of 0 to 10:

- Level of comfort of tactile feedback.
- How instantaneous were the vibrations felt? (short or long and continuous, with higher scores indicating a shorter duration)
- Noticeability of tactile feedback while walking (the higher the score, the more noticeable the feedback).
- How synchronized was the feedback on the opposite arms? (with high scores implying that the feedback felt highly synchronized)
- Intuitiveness of the feedback pattern for adjusting your gait. (with higher scores suggesting the feedback pattern was intuitive)

## III. Results

12 subjects (7 males, 5 females, 24.8 ± 7.0 years with a range of 20-44 years, 69.3 ± 12.4 kg, 1.72 ± 0.12 m, including one subject with self-reported moderate scoliosis) provided informed consent and participated in the study approved by University of Maine’s IRB. To obtain the sagittal angles of the arm and foot segments, the angle between the axes of the IMUs and the local earth-fixed coordinate frame was utilized [38]. To quantify the subjects’ gaits, 5 parameters were calculated; arm cycle time (*T*_*a*_) and foot cycle time (*T*_*f*_) were defined as the time between the consecutive peaks of their sagittal angles, which corresponded to heel strikes and maximum flexion angle for the foot and arm angles, respectively (see Fig. 2). The arm range of motion or amplitude (Θ_*a*_) was calculated as the difference between the maximum and minimum of the arm angle in each gait cycle. Integration of acceleration data and a zero velocity update method [41] were employed to calculate stride length (*L*) and stride velocity (*V*).

In all the one-lap trials, 15 consecutive cycles before the subject started decelerating to stop were used to calculate the parameters of interest for both the left and right sides which were then averaged. For trial *N*, the same 100 cycles for calculating 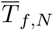 during the experiment were selected. For trial *HM*_0.9_, 15 cycles from when feedback was enabled and 15 cycles after disabling it and towards the end of the trial were compared.

Statistical analyses were performed in SPSS v27 (IBM Corporation, Armonk, NY, USA). When performing repeated measures ANOVA, the sphericity assumption was tested with Mauchly’s test and Greenhouse-Geisser correction was applied if it was rejected. If significant main effects were found, the one-way ANOVA was followed by post-hoc tests with Bonferroni correction to identify the pairs with a significant difference.

### A. Spatiotemporal Parameters and Arm ROM

To determine the effect of haptic feedback on the subjects’ gait, we explored changes in 5 parameters (*T*_*a*_,*T*_*f*_, *L,V*, and Θ_*a*_) in the main feedback trials (*H*_*x*_) and compared them to normal and fast walking conditions and to each other. Fig. 3 demonstrates representative examples of the arm angle during self-selected speeds and two feedback conditions during 10-seconds intervals.

**Fig. 3:**
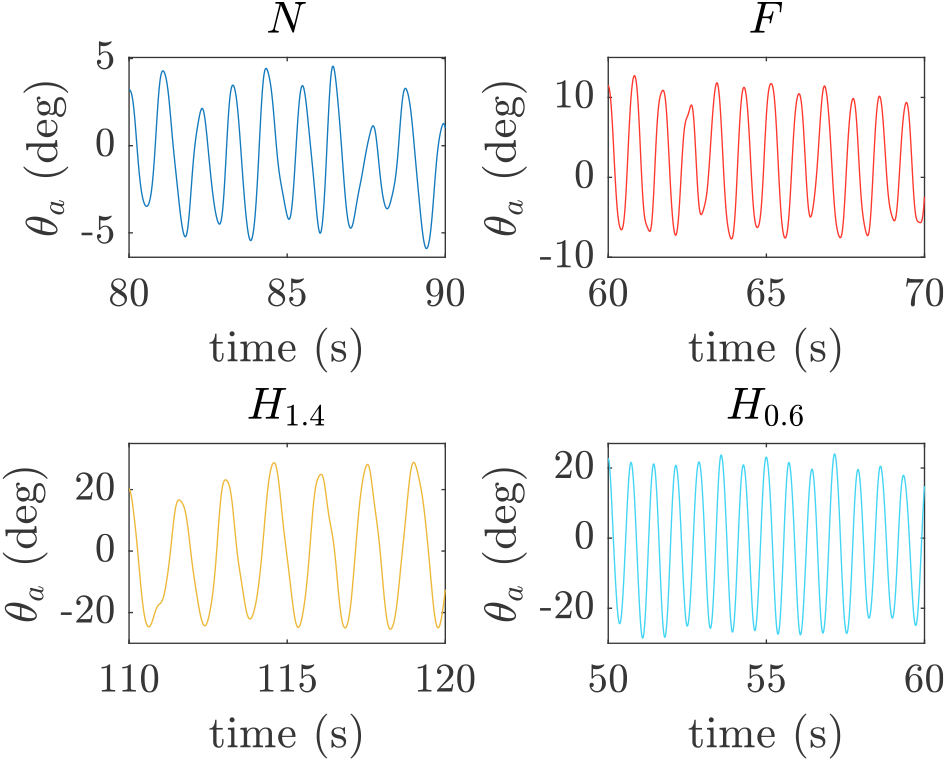
Arm angle (*θ*_*a*_) of one subject shown in four different conditions for 10-seconds intervals. For this figure, to remove the initial offset of *θ*_*a*_, a zero-lag high-pass filter with a cutoff frequency of 0.01 Hz was utilized.

Fig. 4a shows the bar plots for the arm and foot cycles times in different trials. Significant differences were identified across the conditions for both *T*_*a*_ (*F* (1.94, 21.3) = 102.5, *p* < 0.001) and *T*_*f*_ (*F* (2.24, 24.6) = 65.1, *p* < 0.001). The results of the post-hoc tests for *T*_*a*_ are summarized in the figure. In terms of significance, similar results were observed for the post-hoc test of *T*_*f*_, with the exception of no significant difference between *H*_1.0_ and *F* (*p* = 0.054). Performing independent samples t-tests on *T*_*a*_ and *T*_*f*_ in each trial did not find any significant difference between them (*p >* 0.31). Compared to *N*, the arm cycle time (and foot cycle time) was on average 8.7% (9.1%) lower in *F*, 35.5% (29.7%) higher in *H*_1.4_, 16.9% (14.9%) higher in *H*_1.2_, 0.88% (1.2%) lower in *H*_1.0_, 8.8% (8.2%)\ lower in *H*_0.9_, 15.6% (14.6%) lower in *H*_0.8_, and 20.0% (19.7%) lower in *H*_0.6_. To examine the potential impact of following the feedback on gait variability, we calculated the coefficient of variation of stride time 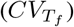 with the results presented in Fig. 4b. Repeated measures ANOVA revealed that there was a significant difference in 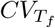 among the trials (*F* (2.06, 22.7) = 0.36, *p* = 0.036) with the results of the post-hoc tests shown in the figure. Compared to normal walking (*N*), the coefficient of variation was on average 43.3% smaller in *F*, 25.1% larger in *H*_1.4_, 0.24% smaller in *H*_1.2_, 35.2% smaller in *H*_1.0_, 39.0% smaller in *H*_0.9_, 16.3% smaller in *H*_0.8_, and 5.30% larger in *H*_0.6_.

**Fig. 4:**
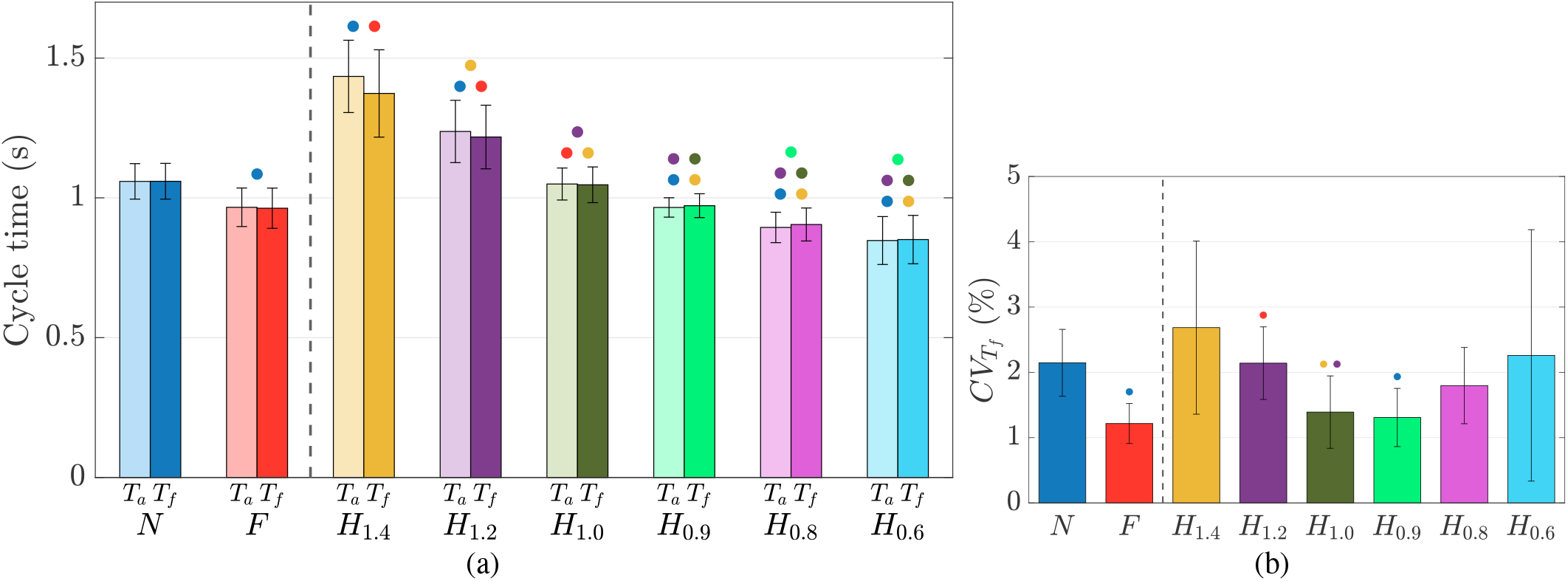
(a) The arm and foot cycle times (*T*_*a*_ and *T*_*f*_) and (b) the coefficient of variation of foot cycle time in different condition. The circles denote significant differences found by the post-hoc tests for *T*_*a*_ (*p* < 0.05) in (a) and for 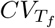 in (b), with the color corresponding to the color of the respective pair’s bar. The dashed lines separate *N* and *F* from the feedback trials.

Fig. 5 shows the arm range of motion (Θ_*a*_) in different conditions. Repeated measures ANOVA found significant differences among the trials (*F* (3.11, 34.2) = 14.5, *p* < 0.001). As shown in the figure, the post-hoc tests showed that Θ_*a*_ was significantly higher in all feedback trials compared to *N* and *F* ; however, this was not the case when comparing *N* and *F* or pairs of feedback trials (*p >* 0.12). Compared to normal walking (*N*), the range of motion was 12% larger in *F* and between 70.9% to 87.8% higher in the feedback trials.

**Fig. 5:**
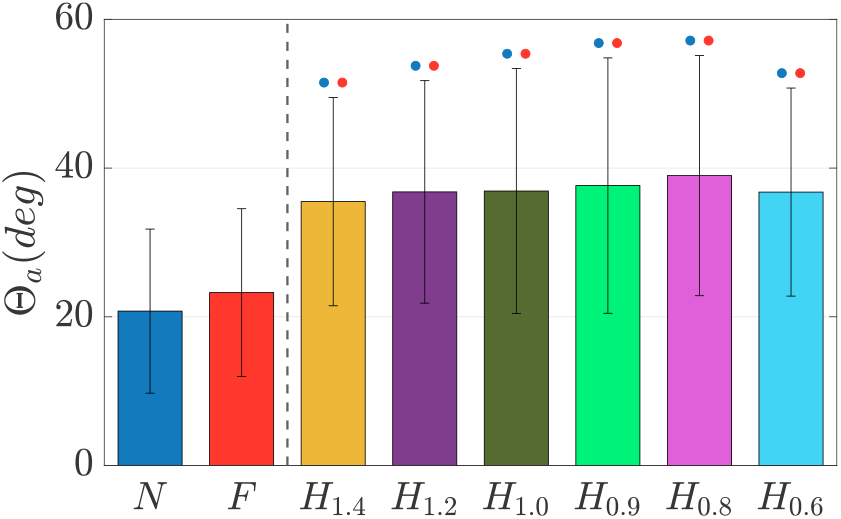
The arm range of motion (arm swing amplitude) in different conditions.

Repeated measures ANOVA of *L* revealed a statistically significant difference for that parameter across the trials (*F* (2.8, 22.9) = 16.0, *p* < 0.001). The values of *L* in different trials and the results are shown in Fig. 6. Compared to *N*, the average stride length was 8.6% larger in *F*, 5.7% smaller in *H*_1.4_, 2.0% smaller in *H*_1.2_, 3.8% larger in *H*_1.0_, 6.3% larger in *H*_0.9_, 7.6% larger in *H*_0.8_, and 8.4% larger in *H*_0.6_.

**Fig. 6:**
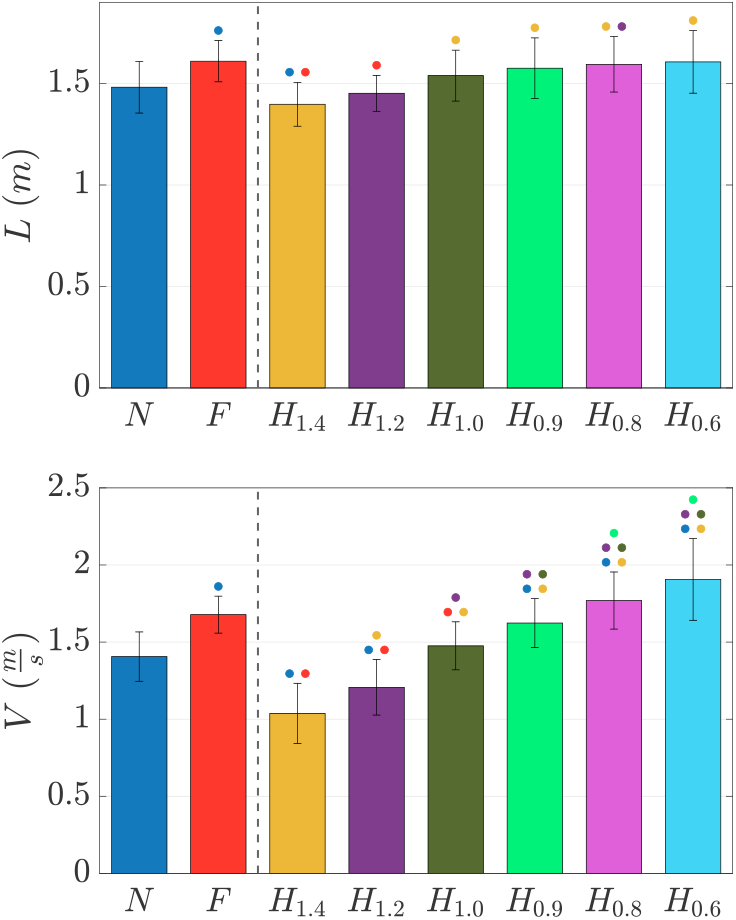
Stride length (top) and stride velocity (bottom) in different trials.

Similar analysis showed significant variations of *V* (*F* (1.86, 20.5) = 53.4, *p* < 0.001) with the post-hoc results shown in Fig. 6. Compared to *N*, the subjects were 19.3% faster in *F*, 26.2% slower in *H*_1.4_, 14.2% slower in *H*_1.2_, 5.0% faster in *H*_1.0_, 15.5% faster in *H*_0.9_, 25.9% faster in *H*_0.8_, and 35.6% faster in *H*_0.6_.

### B. Relationship Between Feedback and Gait Cycle Times

The goal here was to visualize the relationship between the cycle times of the haptic feedback and the cycle time of the user’s gait, which could serve as a baseline for future studies. The independent variable (input) was assumed to be *k*, or the cycle time of feedback relative to 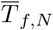. The dependent variable (output) was also a normalized value defined as the ratio of foot cycle time under feedback to 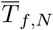.

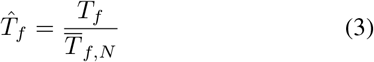

Fig. 7 shows data points and the trends for each individual subject as well as the fitted line (*R*^2^ = 0.80, *p* < 0.001) which has the following equation,

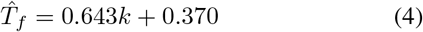

**Fig. 7:**
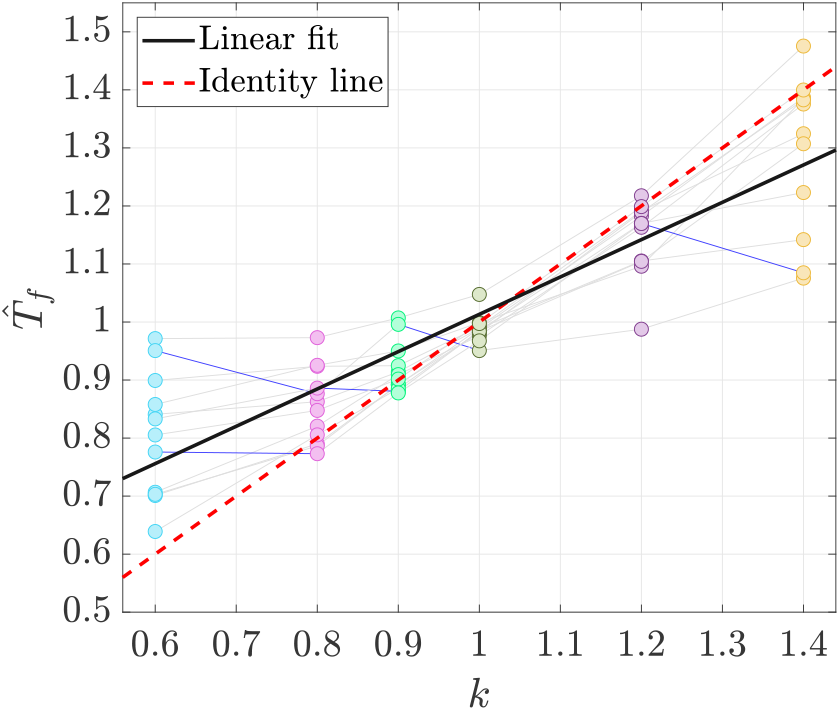
The data points and the linear fit describing the relation between normalized feedback cycle time (*k*) and normalized foot cycle time 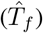. The identity line 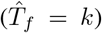 is shown for reference. The blue lines connect that data points of any subject whose cycle times changed conversely to by feedback cycle time (e.g., increased gait cycle time while feedback cycle time decreased), whereas the gray lines show the expected trend.

### C. Transient Response to Feedback

To investigate how quickly the subjects adapted to the feedback and reached a cycle time close to their steady state, we examined the changes in foot cycle time (*T*_*f*_) and arm cycle time (*T*_*a*_) in trials *H*_1.4_ and *H*_0.6_ from the start of the trial (i.e., the transient response). These two trials were selected as they had the slowest and fastest feedback cycle times and thus they were more likely to elicit an immediate response from the subjects. The steady-state values were the 15-cycles averages reported in Fig. 4a and Section III-A, and the end of the transition period was approximated as the first stride that exceeded 98% of the steady-state value, a common criterion for settling time in control engineering. The results are shown in Fig. 8. In both trials, the feedback started after the detection of the 10th heel strike (i.e., at the start of stride 10). In trial *H*_1.4_, the ensemble average of *T*_*f*_ crossed the threshold in 9 strides/8 feedback cycles/12.2 seconds and the ensemble average of *T*_*a*_ in 7 arm movement cycles/6 feedback cycles/10.2 seconds. In trial *H*_0.6_ the threshold was crossed in 5 strides/7 feedback cycles/5.4 seconds and 2 arm movement cycles/3 feedback cycles/2.7 seconds by *T*_*f*_ and *T*_*a*_, respectively.

**Fig. 8:**
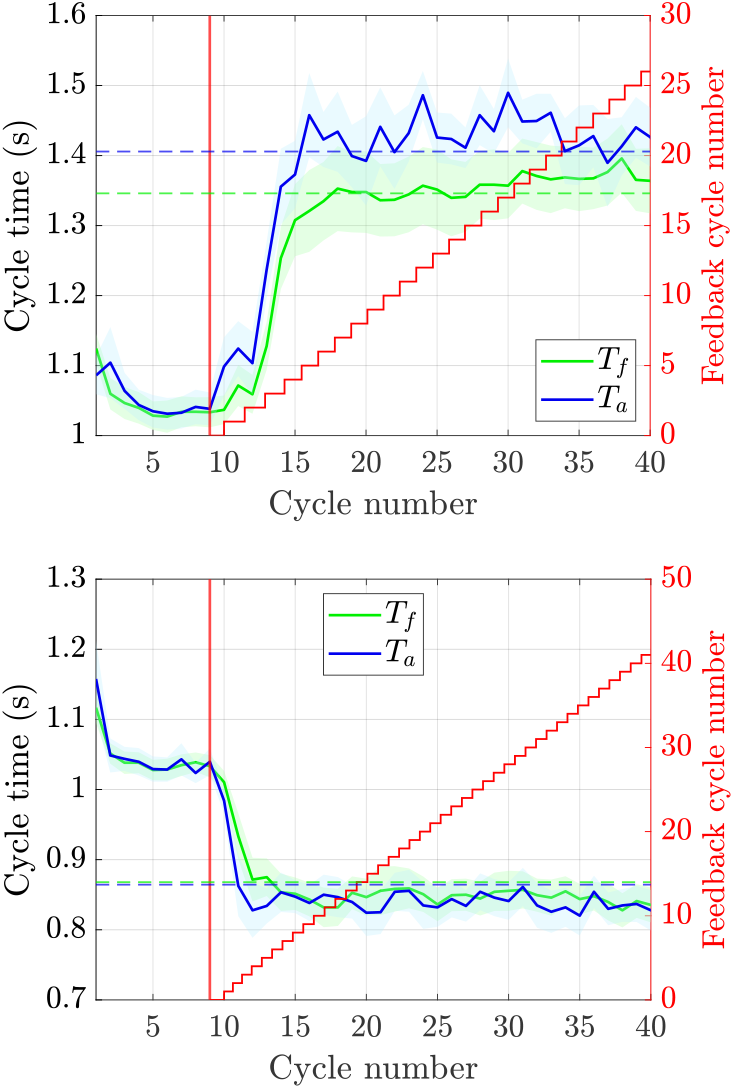
Response of the foot and arm cycle times (*T*_*f*_ and *T*_*a*_, respectively) to the feedback in *H*_1.4_ (top) and *H*_0.6_ (bottom), shown from stride 1 to 40. The solid green and blue lines are the ensemble averages of all subjects and the colored regions show the points within 1 standard error of the averages. The red vertical lines specify the start of feedback and the dashed lines show 98% of the average value of *T*_*f*_ and *T*_*a*_ in the corresponding trial as reported in Fig. 4a. The stair graphs show when feedback cycles started within each stride and the feedback cycle number is shown on the right y-axis.

In addition to ensemble average, we also calculated the transient response for each subject. Here, the parameter of interest was the time (in seconds) between the start of feedback and when *T*_*f*_ and *T*_*a*_ reached their respective steady-state thresholds. In trial *H*_1.4_, this time was approximately 13.3 ± 8.0 seconds for *T*_*f*_ and 8.90 ± 2.9 seconds for *T*_*a*_. In trial *H*_0.6_, the time was calculated to be 4.56 ± 1.6 seconds for *T*_*f*_ and 4.04 ± 1.4 seconds for *T*_*a*_. Paired t-tests for *H*_1.4_ and *H*_0.6_ did not identify a significant difference between *T*_*f*_ and *T*_*a*_ (*p >* 0.31). However, the settling times for both foot and arm in *H*_0.6_ were more similar to each other and faster than the ones in *H*_1.4_.

### D. *Maintaining the Rhythm (HM*_0.9_)

Using paired t-tests, the 5 parameters were compared during the portion of the *HM*_0.9_ when feedback was enabled and with the portion when it was disabled. No significant difference was found for any of the parameters with *p* = 0.807 for *T*_*a*_, *p* = 0.42 for *T*_*f*_, *p* = 0.16 for *L, p* = 0.50 for *V*, and *p* = 0.068 for Θ_*a*_. Similar analysis comparing the parameters when feedback was disabled to *H*_0.9_ found no significant differences between them in the two trials (*p >* 0.2).

### E. *Feedback on One Side Only (HO*_0.9_)

Comparing the 5 parameters on the side that did receive feedback to the other side using paired t-tests, no significant difference between them was identified (*p >* 0.23). To determine whether providing feedback on only one side or both sides created a difference in the overall gait, the results of the feedback and no-feedback sides were averaged and compared with those of trial *H*_0.9_ by performing paired t-tests. No statistically significant difference was observed for any of the parameters (*p* = 0.26 for *T*_*a*_, *p* = 0.52 for *T*_*f*_, *p* = 0.52 for *L, p* = 0.52 for *V*, and *p* = 0.098 for Θ_*a*_). There was an even split in the subjects’ preferences for receiving feedback on one vs. both sides; 6 subjects (50%) found the feedback on one side to be easier to follow while the other 6 subjects preferred receiving feedback on both sides. The “no difference” option was not selected by any participant.

### F. *Providing Information About Feedback Cycle Time (HI*_0.8_)

Paired t-tests were performed to compare the results of trial *HI*_0.8_ with those of trial *H*_0.8_. No significant difference was found in any of the 5 gait parameters (*p >* 0.22).

### G. Subjective Ratings

The questions were rated by the subjects as follows:

- Level of comfort of tactile feedback. (8.50 ± 2.07)
- How instantaneous did the vibrations feel? (7.75 ± 1.49)
- Noticeability of tactile feedback while walking. (7.75 ± 1.96)
- How synchronized was the feedback on the opposite arms? (8.75 ± 1.06)
- Intuitiveness of the feedback pattern for adjusting your gait. (7.92 ± 1.51)

## IV. Discussion

### A. Spatiotemporal Parameters and Arm ROM

The subjects adjusted their arm and leg movements in response to the tactile feedback, resulting in a range of change from +35.5% increase to -20% decrease compared to normal walking for arm swing cycle time and similar values for foot (leg) cycle time. The proposed system is essentially a gait pacemaker that can set the rhythm of subjects’ walking. It should be noted that all although the subjects were instructed to just simply match the feedback and thus remained completely naive to whether they should increase or decrease their arm cycle times, they could successfully adapt their movements based on the feedback. Although not statistically significant, the cadences achieved in *H*_0.8_ and *H*_0.6_ were faster than the fast walking trial; however, the absence of a significant difference between *H*_0.8_ and *H*_0.6_ can indicate a possible point of diminishing returns when reducing the feedback cycle time (*k*). Further, the cycle time in the trial with the same rhythm as normal walking (*H*_1.0_) was similar to *N*, suggesting that the feedback did not cause any appreciable bias towards a faster or slower walking rhythm.

An arm-to-leg cycle time ratio close to 1 was maintained in all the feedback trials. However, in those with a higher feedback cycle time than normal walking (*H*_1.4_ and *H*_1.2_), a trend of desynchronization between arms and legs was observed with the arm cycle time being noticeably larger than the foot cycle time, particularly in *H*_1.4_. Except for the trials with the slowest and fastest feedback rhythms (*H*_1.4_ and *H*_0.6_, respectively), other feedback trials had a similar or lower variability than normal walking with the minimum variability achieved in *H*_0.9_. In trial *H*_1.0_ the subjects achieved a walking rhythm similar to normal walking but with a lower (though statically insignificant) variability suggesting that the feedback would not substantially increase the variability. Given the association between stride time variability and the risk of fall in populations such as older adults [42], this is an important result for future applications of the presented system.

No instruction was given to the subjects regarding increasing their arm range of motion (Θ_*a*_). Even so, Θ_*a*_ increased in all feedback conditions compared to normal and fast walking, achieving its highest value in trial *H*_0.8_. The increase occurred naturally as a response to feedback and was at a similar level across the different trials. This increase could be due to the presence of a parameter with the same value in all feedback conditions such as the duration of feedback (*t*_*on*_). It is notable that there was a small decrease in *H*_0.6_ compared to *H*_0.8_ which could be due to the subjects having to reduce their arm swing amplitude in order to maintain the stability of their body rotational motion at faster walking speed as a common strategy for controlling arm swing during fast walking [4]. The significant changes in arm swing amplitude clearly indicates the ability and efficacy of the feedback to effect change arm swing in the subjects and, thereby, modify their gait. These results show that modulating arm swing cycle time may be an effective method to attain improvements in both arm swing amplitude and frequency, where the spontaneous increase in arm swing range of motion could be beneficial in populations with diminished arm swing such as PD patients.

Regarding the changes in stride length *L*, with the exception of *H*_1.4_ and *H*_1.2_, there was no significant difference between the feedback conditions and normal or fast walking. However, by examining *H*_1.0_ to *H*_0.6_, it is clear the average stride lengths in all 4 feedback conditions were above that of normal walking and even reached values at the level of fast walking in trial *H*_0.6_, but there was no statistically significant difference among those 4 feedback conditions. As expected from the observed trends for stride length and cycle time, stride velocity *V* followed a similar trend to cycle time and decreased and increased depending the direction of feedback. Improving gait speed by training the user to follow faster rhythms could be beneficial for populations such as post-stroke and PD patients, as well as older adults, whose ability to generate rhythmic motion is disrupted and suffer from impaired mobility and low walking speeds.

### B. Relationship Between Feedback and Gait Cycle Times

Although it is ideal to have an identical scaling behaviour between the feedback (desired) cycle time and the subject (output) cycle time with no variability, as the output would exactly follow the changes in input in that case, variations in human movements is expected due to factors such as subject heterogeneity. As a result, as described by Eq. 4, the average change in 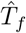 when varying *k* by 10% would be 6.4%.

The linear fit intersected with the identity line at almost *k*=1 as shown in Fig. 7 suggesting the feedback did not create a bias in gait cycle times. The linear fit was below the identity for *k >* 1 (i.e., on average, gait cycle time was faster than feedback in that region) and above the identity for *k* < 1 (i.e., gait cycle time was slower than feedback in that region). Looking at the trend for individual subjects (the grey and blue lines in Fig. 7), 5 instances were identified where 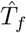 did not decrease with *k*. Out of those 5 (representing 8.3% of total cases), 4 belonged to only 2 subjects. This result demonstrates that generally, despite the subjects being unaware of the timing of the rhythm in each trial, the feedback elicited the expected change in cycle time of each subject across the different trials. It should be noted that there are biomechanical bounds to how much cycle time can be increased or decreased for each individual. Therefore, the relationship between *k* and 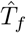 can be best described in full by a nonlinear model in the shape of a possibly asymmetric sigmoid function with an inflection points at *k* = 1. In that case, the biomechanical limits for 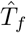 in very high or low values of *k* would manifest in the form of the two asymptotes of the function. For the region of values tested in this experiment, the linear model exhibited good predictive power and therefore could be considered as the linear portion of the sigmoid function that connects the two asymptotic portions. However, there is some saturation effect visible, particularly at *k* = 0.6.

### C. Transient Response to Feedback

Although in this paper we investigated the response of the subjects to a fixed rhythm in order to have a well-controlled experiment, the transient response to the feedback is an important parameter in scenarios where feedback cycle time can vary multiple times as the subject walks. As such, the knowledge of how quickly the subjects adapt to a new rhythm should be incorporated in the design of a trajectory for changing the target cycle time. Our results indicate that it may be necessary to allocate 9 strides (or the equivalent settling time) for allowing the subjects to reach a steady gait after a change of the rhythm. If changing the feedback rhythm during walking is desired, then the system should consider the mentioned settling time and change the rhythm only after the subject has reached the steady-state value. The absence of a noticeable difference between the time it took for the arm and foot cycles times to reach their steady-state values further highlights the presence of interlimb coupling, in which the adaptation of lower limbs’ movements to the arms’ movements occurred quickly and seamlessly.

### D. *Maintaining the Rhythm (HM*_0.9_)

After a short period of receiving feedback, the subjects were able to maintain a gait that was similar to the time when feedback was present. This indicates the potential subjects’ retention of modifications after training with the system, when the feedback is no longer present. Such a retention is critical for training application, where users are expected to attain long-lasting improvements. The effects of longer-term training with the system on users’ gait and retention of improvements require more investigations in the future. Also, the results indicate the possibility of providing a flexible feedback dosage during each training session.

### E. *Feedback on One Side Only (HO*_0.9_)

When feedback was applied to only one side, the subjects were able to maintain a symmetric gait with no significant changes compared to the trial where the tactile feedback was provided to both arms. It is noteworthy that the response to the question immediately after the trial revealed that the subjects did perceive a difference between the two methods (i.e., no subject selected the “no difference option”), but there was a lack of consensus on the preferred strategy. When asked about the reason for their choice, those who preferred feedback on one side provided comments such as “it was easier to focus on one side” and “it was easier to get back on the rhythm after getting out of step with it”, while some in the other group believed that during feedback on one side only “it was easy to forget the other side” and perceived feedback on both sides to “require less information processing”. Therefore, the results demonstrate that, while it is possible to achieve the desired gait by providing feedback to only one arm, the preference of each subject may be taken into account with two short trials at the start of the training session and presenting the two conditions.

### F. *Providing Information About Feedback Cycle Time (HI*_0.8_)

In all the other trials, in order to evaluate the response of the users to only the tactile stimuli, a blind study design was used (i.e., no information regarding the feedback cycle time or rhythm was conveyed to the subjects). In this trial we investigated whether communicating information that exaggerated the feedback cycle time could potentially bias the subject’s perception of the tactile rhythm. The 5 parameters had similar values to the equivalent blind trial (*H*_0.8_) with no significant difference between them, suggesting that the information about feedback can be shared prior to the start of the trial without creating a deviation between the subject’s performance and the expected results.

### G. Subjective Ratings

In the design of the feedback system (Section II-A), we aimed to meet the four requirements of (1) comfort, (2), noticeability, (3) a short perceived duration, and (4) synchronicity of the cues, which we postulate to be vital for a tactile feedback system capable of providing consistent rhythmic cues. The relatively high scores (nearly 8 or above) for all the corresponding questions suggest that the system satisfied those criteria. Further, the subjects found the diagonal feedback scheme intuitive, which is also supported by the fact that after a short familiarization, they were able to adjust their gait accordingly in the trials that followed.

## V. Conclusion

In this work, we presented a wireless and wearable feedback system capable of providing highly synchronized vibrotactile cues. Given the importance of inclusion of arm swing in gait rehabilitation due to interlimb neural coupling, the effect of providing rhythmic tactile feedback on the arms was studied in a variety of conditions in a subject study. The presented system and approach essentially act as a metronome for gait with the vital advantage of engaging both arms and legs in training to promote interlimb coordination and enable neurorehabilitation. With minimal exposure to the system and no information about the frequency of the feedback, the participants could conveniently adjust their gait according to the rhythm of the cues, resulting in reductions of stride time and improvements in stride velocity and arm range of motion. The relationship between the feedback cycle time (input) and the gait cycle time (output) was quantified and the subjects’ transient response to a step input of the feedback was studied. Quantifying the settling times of the arm and foot cycle times in response to step inputs provided a quantitative insight from a dynamic systems viewpoint into arm swing response to tactile stimuli and interlimb coupling. We demonstrated that the subjects were able to retain the changes in their gait after the feedback stopped, could adjust their gait even with feedback applied to only one side, and were not biased by the knowledge about the desired cycle time. The subjective ratings on factors such as noticeability and the perceived synchrony of the cues affirmed our design choices. Future work will include evaluating the system on populations such as older adults and stroke patients, investigating the dose-response effect of training with the tactile cues, studying the upper and lower limb coordination in more detail, and designing closed-loop strategies where the subject’s performance can affect the cues in real-time.

